# Acidic pH is a Metabolic Switch for 2-Hydroxyglutarate Generation and Signaling

**DOI:** 10.1101/051599

**Authors:** Sergiy M. Nadtochiy, Xenia Schafer, Dragony Fu, Keith Nehrke, Joshua Munger, Paul S. Brookes

**Author notes:** To whom correspondence should be addressed: Paul S. Brookes, PhD., Department of Anesthesiology, University of Rochester Medical Center, 601 Elmwood Avenue, Rochester, NY 14642, USA.

## Abstract

2-hydroxyglutarate (2-HG) is an important epigenetic regulator, with potential roles in cancer and stem cell biology. The D (R) enantiomer (D-2-HG) is an *oncometabolite* generated from αketoglutarate (α-KG) by mutant isocitrate dehydrogenase (ICDH), while L (S) 2-HG is generated by lactate dehydrogenase (LDH) and malate dehydrogenase (MDH) in response to hypoxia. Since acidic pH is a common feature of hypoxia, as well as tumor and stem cell microenvironments, we hypothesized that pH may regulate cellular 2-HG levels. Herein we report that cytosolic acidification under normoxia moderately elevated 2-HG in cells, and boosting endogenous substrate α-KG levels further stimulated this elevation. Studies with isolated LDH-1 and MDH-2 revealed that generation of 2-HG by both enzymes was stimulated several-fold at acidic pH, relative to normal physiologic pH. In addition, acidic pH was found to inhibit the activity of the mitochondrial L-2-HG removal enzyme L-2-HG dehydrogenase, and to stimulate the reverse reaction of ICDH (carboxylation of αKG to isocitrate). Furthermore, since acidic pH is known to stabilize hypoxia-inducible factor (HIF), and 2-HG is a known inhibitor of HIF prolyl hydroxylases, we hypothesized that 2-HG may be required for acid-induced HIF stabilization. Accordingly, cells stably over-expressing L-2HGDH exhibited a blunted HIF response to acid. Together these results suggest that acidosis is an important and previously overlooked regulator of 2-HG accumulation and other oncometabolic events, with implications for HIF signaling.

---

The field of cancer biology has long been rapt by the notion of a cancer-specific metabolic phenotype, perhaps most famously embodied in the “Warburg effect”, wherein glycolytic metabolism predominates in cancer cells despite O_2_ availability and largely intact mitochondrial respiratory function (1).

A prominent feature of the cancer metabolic phenotype (reviewed in (2)) is an elevated level of the small metabolic acid 2-hydroxyglutarate (2-HG), derived from the TCA cycle intermediate α-ketoglutarate (α-KG) (3). The D (R) enantiomer of 2-HG (D-2-HG) was shown to be generated by mutant forms of isocitrate dehydrogenase (ICDH) that are associated with a variety of cancers including aggressive gliomas (4). In addition, more recently the L (S) enantiomer (L-2-HG) was shown to be generated under hypoxic conditions by lactate dehydrogenase (LDH) and malate dehydrogenase (MDH) (5, 6). We have also reported elevated D/L-2-HG levels in the heart following ischemic preconditioning (7).

In addition to synthesis, 2-HG levels are regulated by a pair of dehydrogenases that convert 2-HG back to α-KG (i.e., L-2-HGDH and D-2-HGDH). Mutations in these enzymes manifest as the hydroxyglutaric acidurias, devastating inherited metabolic diseases with symptoms including epilepsy and cerebellar ataxia (8-11). However, the importance of these dehydrogenases in regulating 2-HG levels in other settings is not clear.

The downstream signaling roles of D-2-HG in cancer biology and of L-2-HG in hypoxia or stem cell biology, are thought to be mediated by epigenetic effects (12-14), owing to competitive inhibition of the α-KG-dependent dioxygenase superfamily of enzymes. This includes the JmjC domain-containing histone demethylases, the TET 5-methylcytosine hydroxylases, the EGLN prolylhydroxylases that regulate hypoxia inducible factor (HIF), and the AlkB homolog family of DNA/RNA demethylases (15-18). As such, 2-HG is a potentially important link between metabolism and epigenetic signaling.

A common feature of hypoxia, as well as the tumor and stem cell microenvironments, is metabolic acidosis. However, the role of pH in regulating 2-HG formation and disposal has not been considered. We tested the hypothesis that acidic pH is a regulator of 2-HG metabolism at the cellular, mitochondrial, and isolated enzyme levels. Our results indicate that acidic pH can independently drive elevated 2-HG levels, and we propose that pH regulation of 2-HG may have important implications for 2-HG signaling in hypoxia and other settings.

## RESULTS & DISCUSSION

We investigated the role of pH as an independent variable in the regulation of 2-HG levels in HEK293 cells, using an ammonium plus 5-(N-Ethyl-N-isopropyl) amiloride (EIPA) pH clamp system (19). This system was capable of depressing and maintaining intracellular pH to 6.8 within 2 hrs. (Figures 1A–C). This magnitude of pH depression (approximately 0.5 units) is similar to that previously reported upon exposure of cells to hypoxia/anoxia (20-22). As such, the pH-dependent signaling effects measured herein can be considered to represent those that may be expected to occur in hypoxic tissues.

**Figure 1.**
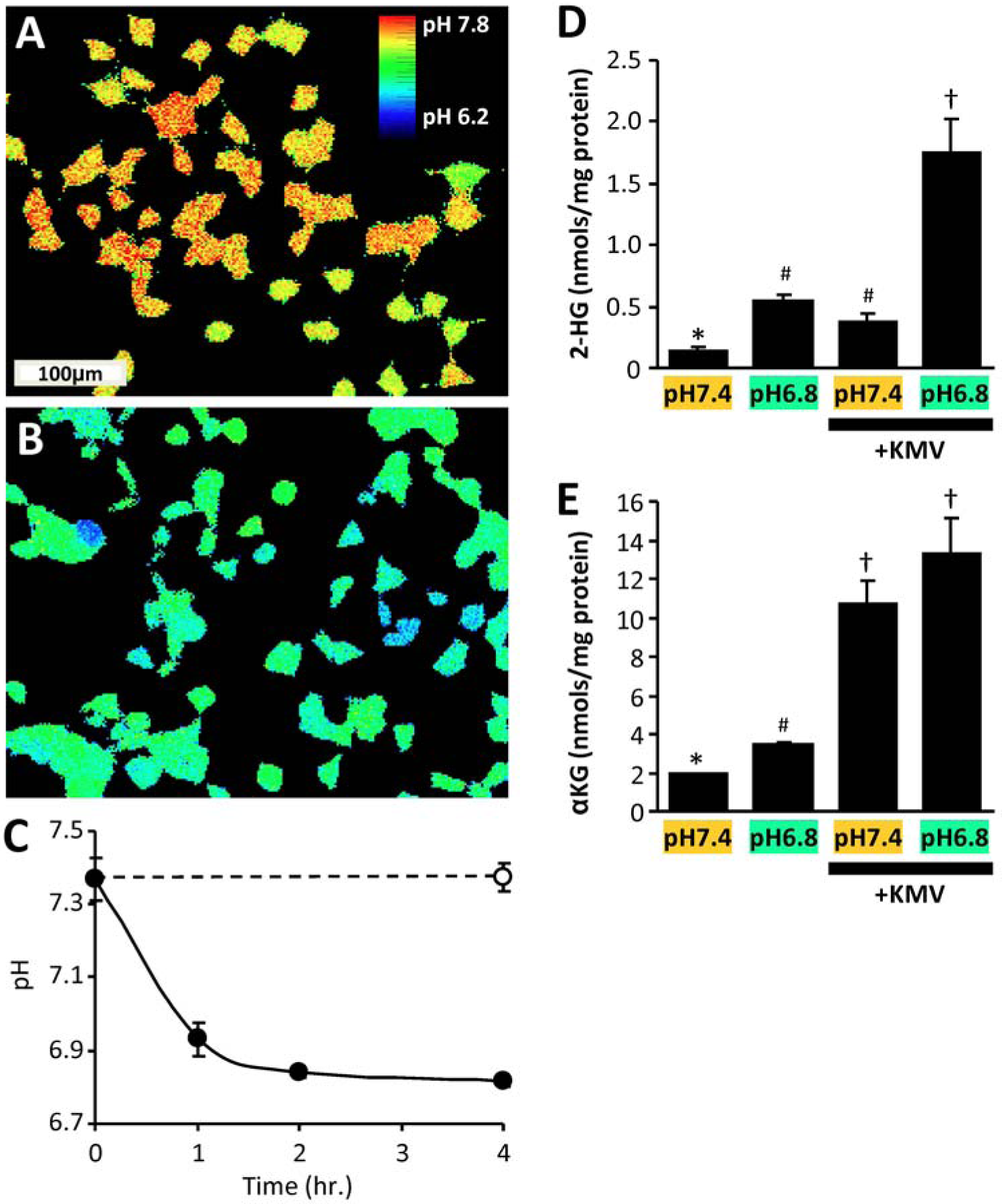
Acidic pH drives 2-HG production in cells. HEK293 cells were maintained at pH 7.4, or incubated under conditions to acidify the cytosol (see methods). **(A/B):** Pseudocolored 490nm/440nm dual excitation ratiometric images of BCECF fluorescence, showing intracellular pH for cells maintained at pH 7.4 (panel A), or pH 6.5 with EIPA (panel B). **(C):** Quantitative time course of cellular acidification. Data are means ± SD, N=3. **(D):** Cellular 2-HG levels measured by LC-MS/MS following incubation at different cellular pH levels in the absence or presence of the α-KGDH inhibitor KMV. **(E):** Cellular α-KG levels measured by LC-MS/MS following incubation at different cellular pH levels in the absence or presence of KMV. Data are means ± SEM, N=6. Different symbols above bars (* ^#^ †) indicate statistically significant differences (two-way ANOVA and Student’s t-test) between groups – i.e. bars with the same symbol are not different.

Although this degree of acidification drove a 4-fold elevation in 2-HG^1^ levels (Figure 1D, left two bars), this was somewhat lower than the >8-fold elevation reported in the same cell type in response to hypoxia (5, 6). Since 2-HG generation also depends on the availability of substrate α-KG, which is elevated in hypoxia (6), we hypothesized that α-KG may be limiting in acidic pH alone. As such, we found that inhibiting the TCA cycle enzyme α-KGDH with ketomethyvalerate (KMV) boosted cellular α-KG levels by ~5-fold regardless of pH (Figure 1E). Furthermore, while KMV alone drove a small increase in 2-HG levels at pH 7.4 (Figure 1D), the combination of KMV plus acidic pH yielded an overall 13-fold increase in 2-HG levels. Together these data indicate that reproducing the acidic pH and elevated α-KG levels seen in hypoxia is sufficient to increase 2-HG in cells. Acidosis may therefore be an important variable that drives 2-HG elevation in hypoxia.

Operating on the premise that the 2-HG observed in cells was the L-isomer (see footnote^1^), we next investigated the mechanism of acid-induced 2-HG elevation, by examining the effects of pH on the known enzymes that generate and remove this metabolite – namely LDH, MDH, ICDH and 2-HGDH. The purity of commercial isolated LDH-1, MDH-2 and NADP^+^-dependent ICDH were verified by gel electrophoresis, with ICDH found to be somewhat impure (Figure 2A). A spectrophotometric assay for the consumption of NADH by LDH in the presence of α-KG (Figure 3A inset) was performed across a range of pH values spanning 6.6–7.8 (i.e. encompassing the pH values studied in intact cells), and revealed that NADH consumption was approximately 2.5-fold faster at pH 6.8 than at pH 7.4 (Figure 3A). In agreement with the cell data (Figure 1D), the rate was also [α-KG] dependent, being faster at 10 mM than at 1 mM α-KG. The spectrophotometric data were confirmed by LC-MS/MS measurement of 2-HG generation (Figure 3B), with approximate molar equivalents of 2-HG produced across the pH range 6.6–7.8 (Figure 2B, r^2^ = 0.97).

**Figure 2.**
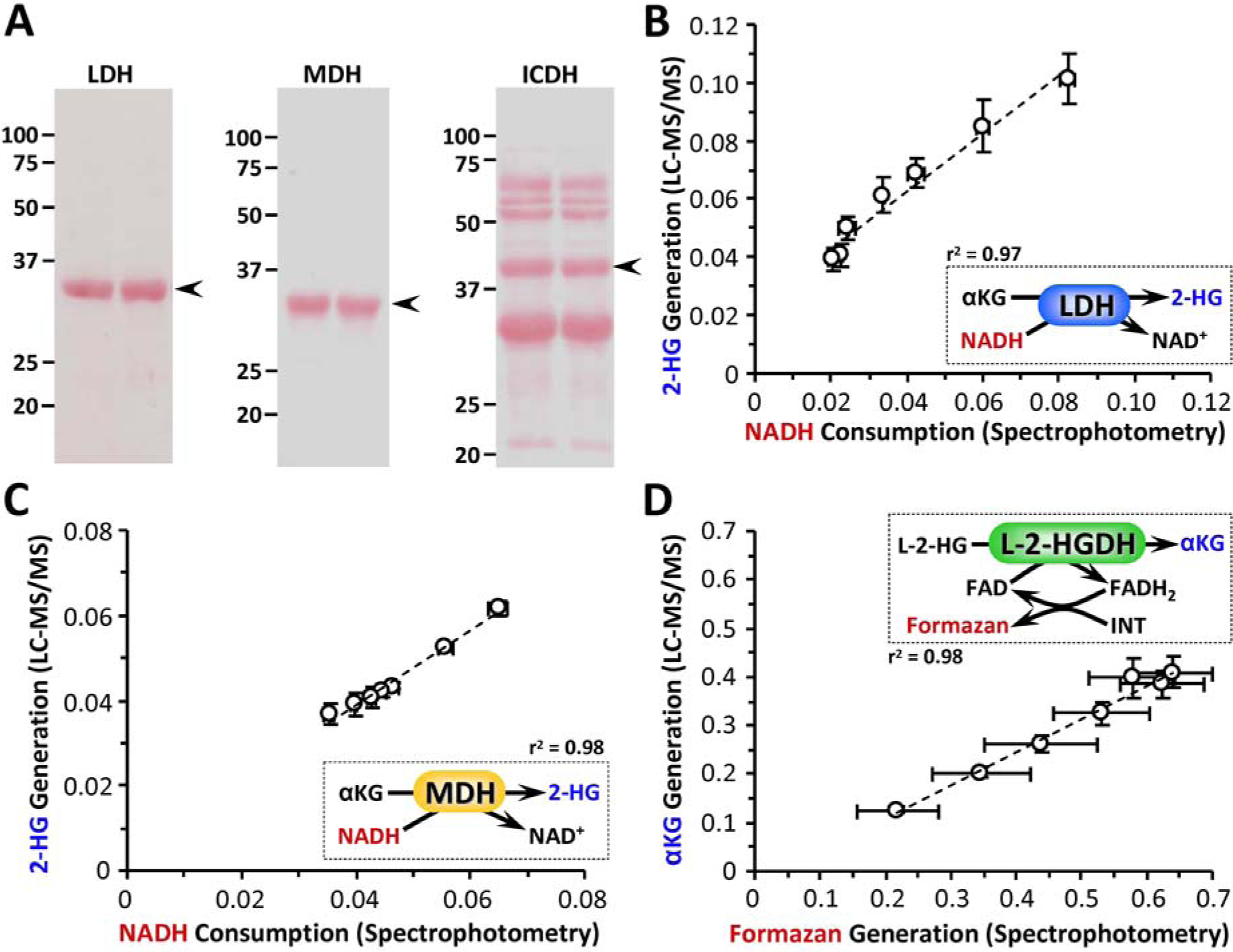
Enzyme purity and Assay Validation. **(A):** SDS-PAGE analysis of commercially obtained LDH (Sigma L3916), MDH (Sigma M2634) and ICDH (Sigma #I2002). Following electrophoresis on 12.5% SDS-PAGE, gels were transferred to nitrocellulose membranes and stained with Ponceau S. Numbers to the left of images show molecular weight markers, in kDa. Data are shown for 2 separate batches of material (2 lanes). Arrows to the right of images show expected position of band, for published molecular weight of the protein. **(B):** Comparison of NADH spectrophotometric assay vs. 2-HG LCMS/MS assay, for the 2-HG generating activity of LDH (i.e., data from Figures 3A and 3B of main manuscript). Reaction is shown in the inset, with red/blue colored parameters matching appropriate axes on the graph. Data are means ± SEM. Dotted line shows linear regression curve fit with correlation coefficient shown alongside inset. **(C):** Comparison of NADH spectrophotometric assay vs. 2-HG LCMS/MS assay, for the 2-HG generating activity of MDH (i.e., data from Figures 3D and 3E of main manuscript). Reaction is shown in the inset, with red/blue colored parameters matching appropriate axes on the graph. Data are means ± SEM. Dotted line shows linear regression curve fit with correlation coefficient shown alongside inset. **(D):** Comparison of Formazan spectrophotometric assay vs. α-KG LCMS/MS assay, for the L-2-HG consuming activity of L-2-HGDH (i.e., data from Figures 6A and 6B of main manuscript). Reaction is shown in the inset, with red/blue colored parameters matching appropriate axes on the graph. Data are means ± SEM, with N for each measure corresponding to the parent data sets in Figures 2 and 3 of the main manuscript. Dotted line shows linear regression curve fit with correlation coefficient shown alongside inset.

The native activity of LDH (i.e., conversion of pyruvate to lactate) was unaffected by pH (Figure 3C); unsurprising for an enzyme that typically drives metabolic acidosis. Fractionally, the 2-HG generating capacity of LDH was at least 4 orders of magnitude lower than its native activity (Figure 3A vs. 3C), indicating that acidic pH unmasks a latent 2-HG generating activity in LDH without impacting its native activity. This observation also suggests that the generation of 2-HG is not a quantitatively important metabolic sink for NADH.

**Figure 3.**
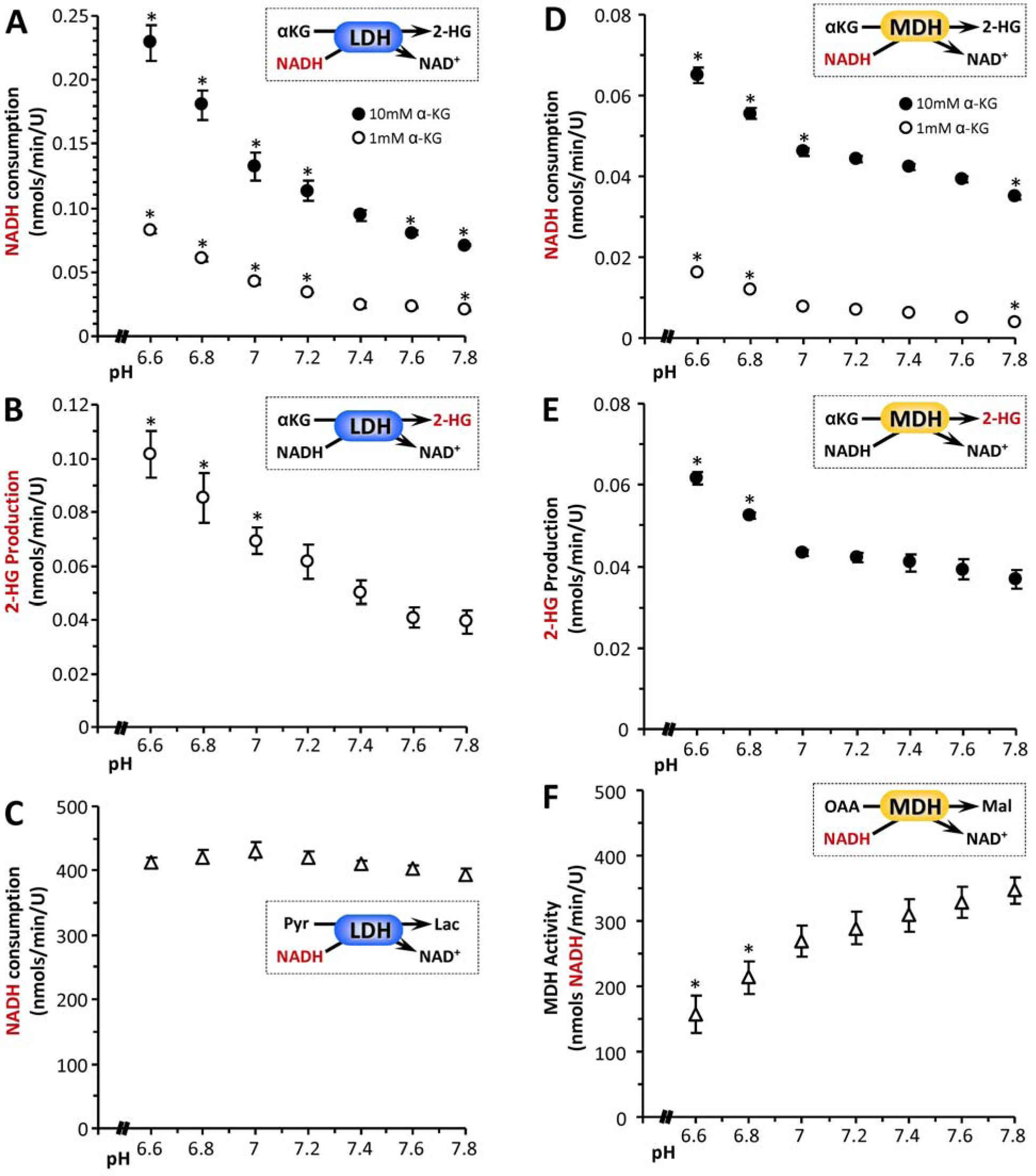
pH dependency of 2-HG generation by lactic or malic dehydrogenases. The 2-HG synthetic activity of isolated LDH was assayed spectrophotometrically as NADH consumption (panel A) or by direct LC-MS/MS assay of 2-HG formation (panel B). Insets to each panel show reaction schemes, with the measured parameter (y-axis of graph) highlighted in red. **(A):** NADH consumption at various pH values, for LDH in the presence of 1 mM (white circles) or 10 mM (black circles) α-KG. **(B):** 2-HG production at various pH values, for LDH in the presence of 1 mM α-KG. **(C):** Native pyruvate to lactate converting activity of LDH at various pH values, assayed spectrophotometrically as NADH consumption. Similar experiments were also performed with isolated MDH. **(D):** NADH consumption at various pH values, for MDH in the presence of 1 mM (white symbols) or 10 mM (black symbols) α-KG. **(E):** 2-HG production at various pH values, for MDH in the presence of 10 mM α-KG. (Note – with MDH, the high concentration of α-KG was necessary due to the low sensitivity of LC-MS/MS based 2-HG assay). **(F):** Native MDH activity (oxaloacetate to malate) at various pH values, assayed spectrophotometrically as NADH consumption. (Note – MDH activity was measured in the reverse direction, i.e., oxaloacetate to malate, due to thermodynamic constraints as detailed in the supplemental methods). All data are means ± SEM, N>3. *p<0.05 (two-way ANOVA followed by Student’s t-test) compared to corresponding value at pH 7.4. Where error bars appear absent, they are sufficiently small as to be wholly contained within the data symbols.

As a control, the effect of hypoxia on the ability of isolated LDH to generate 2-HG at pH 7.4 was also studied. Hypoxia was verified by measuring the spectrum of oxy- vs. deoxy-hemoglobin in the cuvet (Figure 4A). In contrast to the situation seen in cells (5, 6), no effect of pO_2_ on 2-HG generation by isolated LDH was seen (Figures 4B and 4C). These data suggest that hypoxic stimulation of 2-HG generation by LDH is not a property of the isolated enzyme, but rather it requires the intact hypoxic cell environment, likely including acidosis.

**Figure 4.**
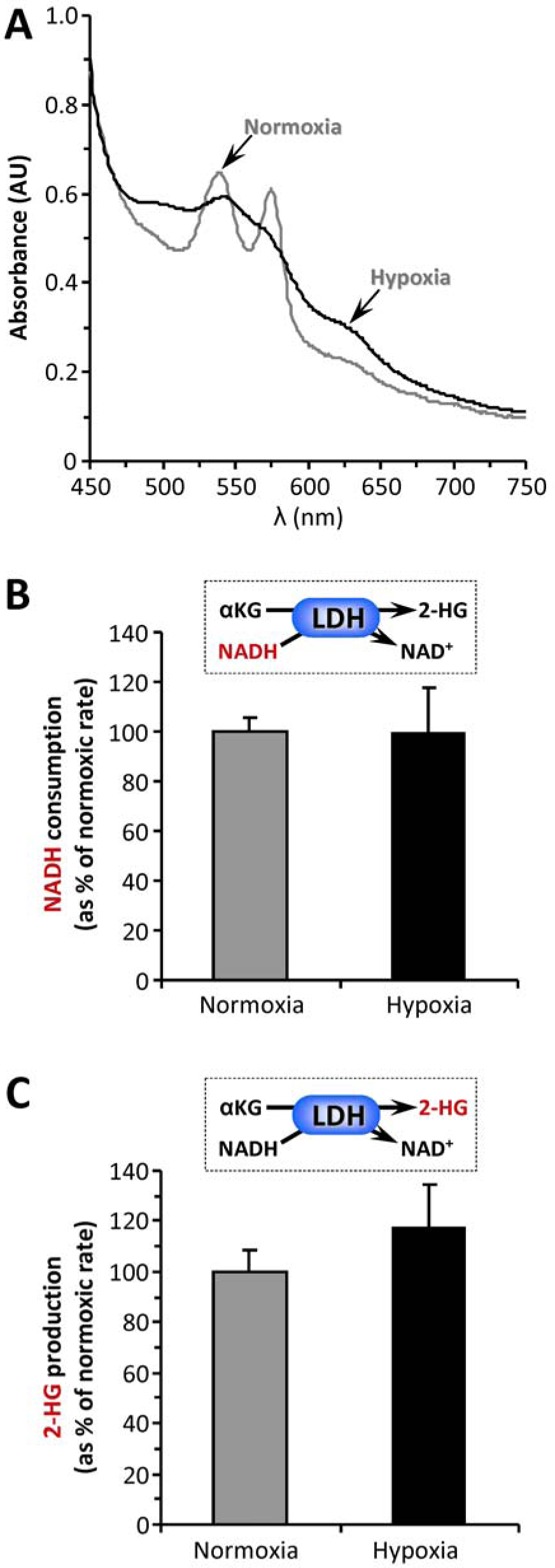
Effect of Hypoxia on 2-HG Generation by LDH. Isolated LDH was incubated in an open flow spectrophotometer cell, as detailed in the methods. A parallel incubation contained 100 μM hemoglobin, to confirm deoxygenation of the sample. **(A):** Absorbance spectra of purified Hb under normoxic (gray line) or hypoxic (black line) conditions, the latter comprising a 1 hr. of argon purge. **(B):** NADH consumption by LDH under normoxic or hypoxic conditions. Data are expressed normalized to the normoxia values, and are means ± SEM. **(C):** 2-HG generation by LDH under normoxic or hypoxic conditions. Samples were removed from the same incubations as panel B and rapidly deproteinized for LC-MS/MS analysis of 2-HG levels. Data are expressed normalized to the normoxia values, and are means ± SEM.

Similar to LDH, the generation of 2-HG by MDH was also found to be higher at pH 6.8 than at pH 7.4 (Figures 3D and 3E). As seen with LDH, spectrophotometric and LC-MS/MS data for MDH correlated well (Figure 2C, r^2^ = 0.98), with approximate molar equivalents of 2-HG produced across the pH range 6.6–7.8. Notably, unlike LDH (Figure 3C) the native activity of MDH was itself pH sensitive, being inhibited ~30% at pH 6.8 versus pH 7.4 (Figure 3F). At first glance, this result might suggest an acid-induced activity switch, away from native and toward 2-HG-generation. However, it should again be noted that the 2-HG generating activity of MDH was at least 3 orders of magnitude lower than its native activity. Although 2-HG generation has been termed a “metabolic error” and the 2-HG dehydrogenases referred to as metabolite repair enzymes (11), our data suggest that the 2-HG synthetic capacity of MDH unmasked by acidic pH is fractionally small, and does not represent a major diversion of carbon away from the TCA cycle.

Comparing LDH and MDH, at the pH range seen in cells (6.8 vs. 7.4), and at the same α-KG level (10 mM), the relative acid-induced stimulation of 2-HG generation was slightly larger for LDH (2-fold) vs. MDH (1.3-fold). In addition, the pH response of LDH appeared to initiate at slightly more neutral pH, whereas the MDH response was flatter until more acidic pH values were attained. These data, plus the >3-fold greater specific activity for 2-HG generation by LDH vs. MDH, suggest that LDH may be a quantitatively more important source of 2-HG in response to acid.

The ability of native (non-mutant) ICDH to convert α-KG to 2-HG in a pH-sensitive manner was also tested. While ICDH in the presence of αKG indeed consumed NADPH in a manner that was accelerated at acidic pH (Figure 5A), no 2-HG generation was seen (Figure 5B). Instead, ICDH was observed to carboxylate α-KG to isocitrate, and this reaction was stimulated at acidic pH (Figure 5C). This is somewhat expected because the reaction uses protons as a substrate (Figure 5C inset). Such reversal of ICDH activity was previously shown to be triggered under hypoxic conditions (23), and is an important pathway for diversion of carbon from anaplerotic glutamine toward lipid synthesis (via ATP citrate lyase) in cancer cells. Thus, while native ICDH does not appear to be a source for acid-triggered 2-HG generation, these data suggest that acidic pH may regulate numerous oncometabolic pathways, including reductive α-KG carboxylation.

**Figure 5.**
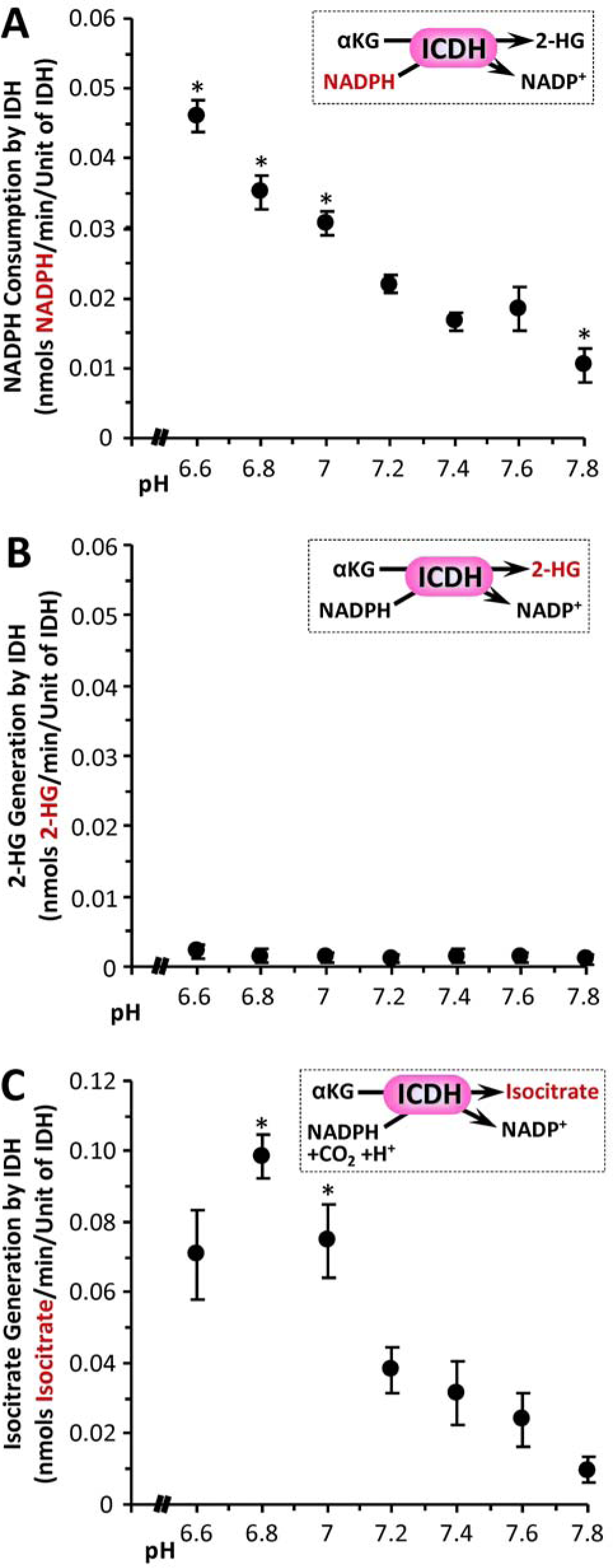
pH dependency of isocitrate dehydrogenase activity. The ability of commercial native ICDH (Figure 2A) to catalyze the generation of 2-HG from α-KG was monitored spectrophotometrically as NADPH consumption (panel A) or by direct LC-MS/MS assay of 2-HG formation (panel B). Insets to each panel show reaction schemes, with the measured parameter (y-axis of graph) highlighted in red. **(A):** NADPH consumption at various pH values, for ICDH in the presence of 10 mM α-KG. **(B):** 2-HG production at various pH values, for ICDH in the presence of 10 mM α-KG. Although NADPH was consumed, 2-HG was not generated. **(C):** Reverse activity of ICDH to generate isocitrate from α-KG, monitored by LC-MS/MS assay of isocitrate formation (see panel inset for reaction scheme). Data are means ± SEM. N=7-8. *p<0.05 (two-way ANOVA followed by Student’s t-test) compared to corresponding value at pH 7.4.

Since cellular 2-HG levels are also regulated by specific dehydrogenases, we next tested the effects of pH on mitochondrial L-2-HGDH activity, using both a formazan-linked colorimetric assay (10) and LC-MS/MS to quantify α-KG generation from the substrate L-2-HG. Both assays correlated well (Figure 2D, r^2^ = 0.98), and showed that L-2-HGDH activity was inhibited by acidic pH (Figure 6). Together with the data in Figure 3 this result indicates that acidic pH both promotes the formation of L-2-HG and inhibits its removal, suggesting a coordinated metabolic response to elevate L-2-HG under acidic conditions. Based on the data in Figures 1, 3 and 6, we conclude that pH is an important determinant of cellular 2-HG levels, and that acidosis may play a key role in the L-2-HG elevation observed in hypoxia (5, 6).

**Figure 6.**
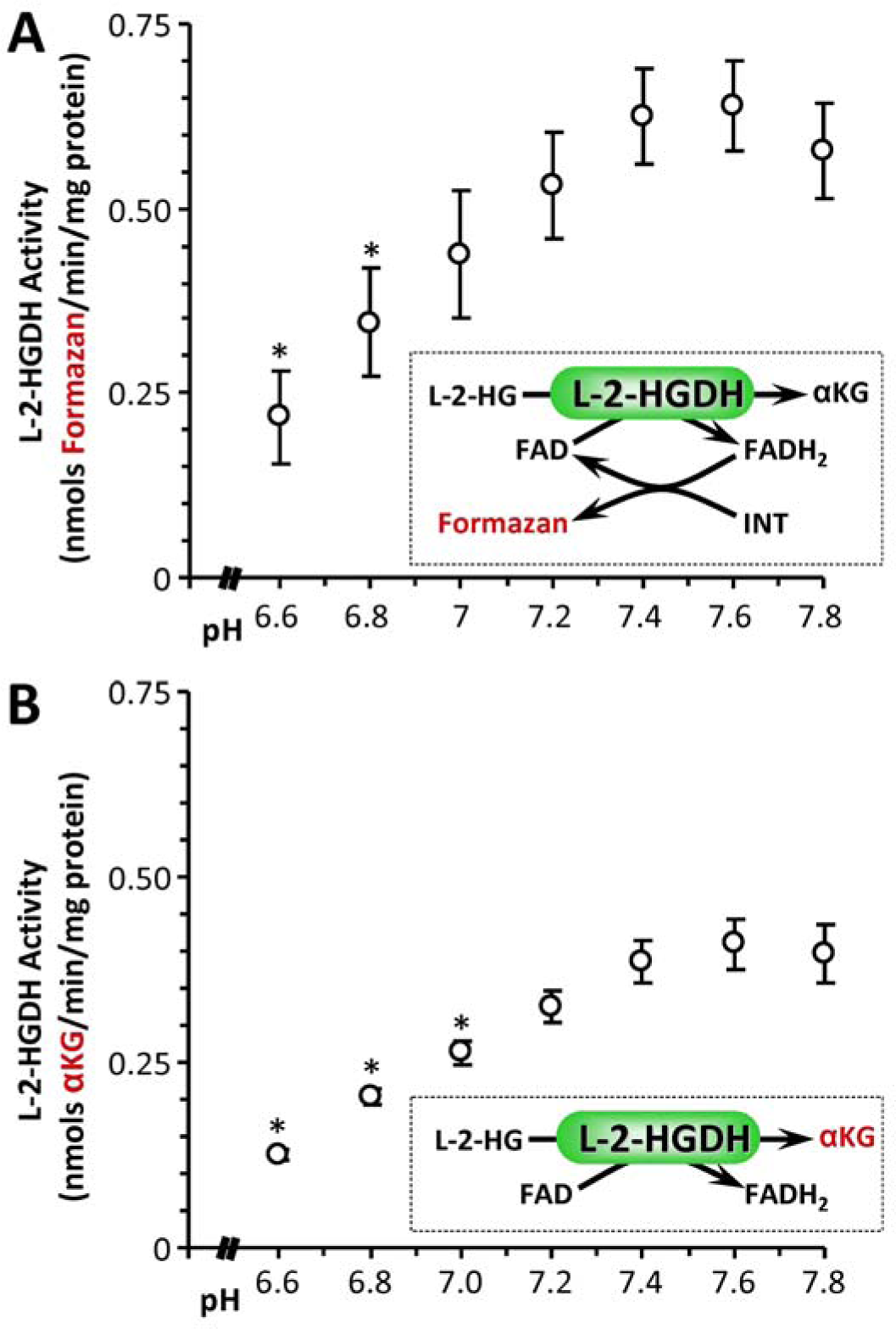
pH dependency of L-2-HG dehydrogenase (L-2-HGDH). The L-2-HGDH activity of isolated permeabilized mouse heart mitochondria was measured spectrophotometrically using either an iodonitrotetrazolium (INT) to formazan coupled assay (panel A) or by direct LC-MS/MS assay of α-KG formation (panel B). **(A):** Formazan production at various pH values, for mitochondria supplied with 500 μM L-2-HG. **(B):** α-KG production at various pH values, for mitochondria supplied with 500 μM L-2-HG. Data are means ± SEM, N=4. *p<0.05 (two-way ANOVA followed by Student’s t-test) compared to corresponding value at pH 7.4.

Biochemically, the reaction catalyzed by LDH and MDH during generation of 2-HG is the same as the native reactions of these enzymes, namely the NADH dependent conversion of an αketo acid to its corresponding α-hydroxy acid (Figure 7). The only difference between substrates is the corresponding R group, and our current findings suggest that the mechanism of acid-induced 2-HG generation by LDH and MDH may involve changes in access to the active site, permitting entry of a larger R group. Such a model leads to a simple prediction: if LDH under acidic conditions can accommodate the R group of α-KG ((CH_2_)_2_–COOH)), it should also be able to accommodate oxaloacetate (OAA). Figures 7B and 7C show that this prediction is correct, with LDH able to convert OAA to malate (i.e. to perform the MDH reaction) in an acid-sensitive manner. A crystal structure for LDH in the presence of the inhibitor oxamate (R group = NH_2_, PDB 1I0Z), shows the region of the protein that may accommodate the R group occupied by 2 molecules of water, with no acid-sensitive residues such as histidine nearby (24). As such, the mechanism by which acidic pH may allow larger substrates into the active site of LDH remains to be elucidated. In addition, the ability of LDH (or MDH) to accept even larger substrates such as αketobutyrate is unknown, but could be worthy of investigation because elevated α-hydroxybutyrate has been shown to be a component of the metabolic signature of mitochondrial dysfunction across multiple organisms (25, 26). In general, the potential for acidic pH to remodel metabolite profiles by promoting α-keto to α-hydroxy acid conversion, remains largely unexplored.

**Figure 7.**
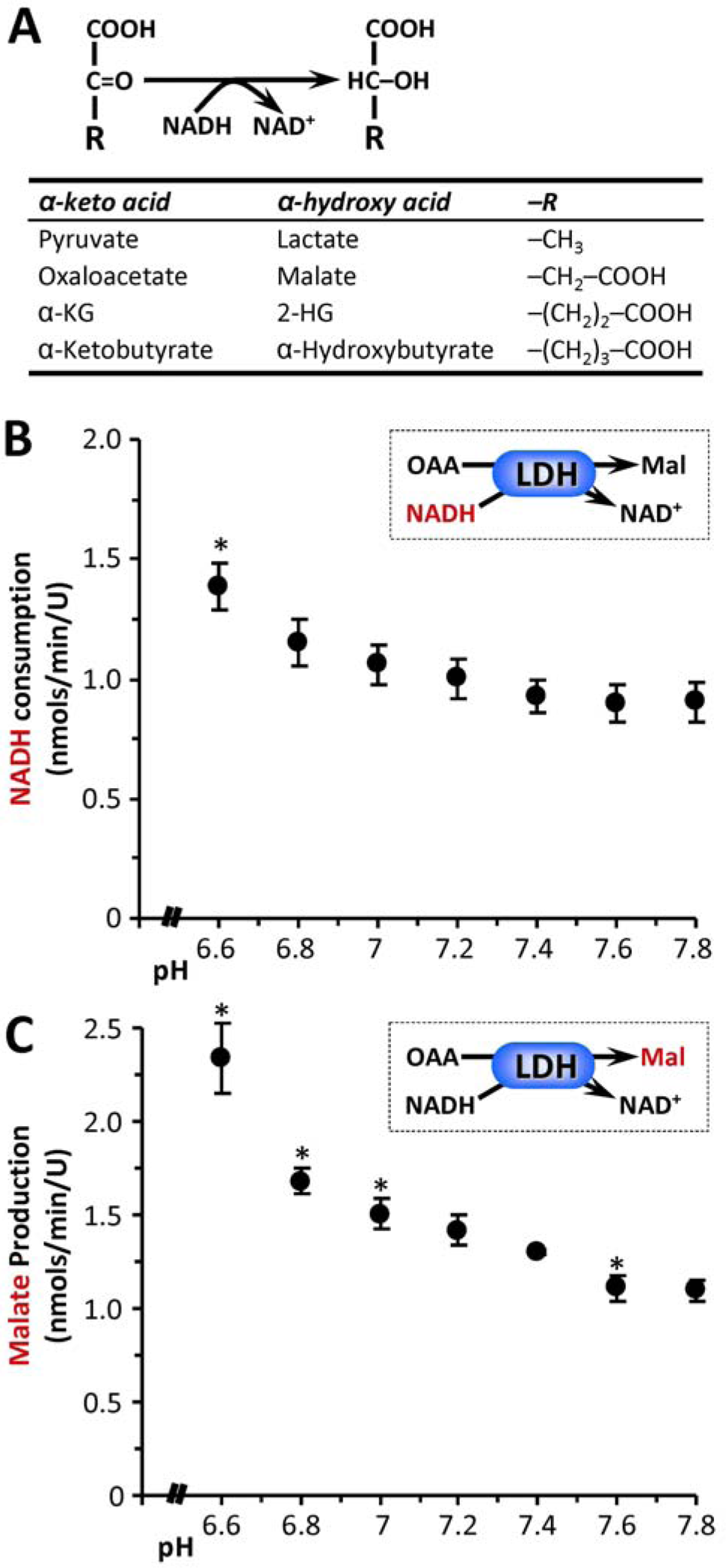
Acid induced substrate promiscuity in α-keto acid dehydrogenases. **(A):** Generalized reaction scheme for α-keto acid dehydrogenases, with the table showing specific examples and the corresponding –R group for each substrate/product pair. **(B):** The ability of LDH to catalyze the native back reaction of MDH (i.e. OAA to malate) was measured at various pH values, assayed spectrophotometrically as NADH consumption. **(C):** Generation of malate in this system was also assayed by LC-MS/MS. Data are means ± SEM, N=4. *p<0.05 (two-way ANOVA followed by Student’s t-test) compared to corresponding value at pH 7.4.

Downstream signaling by 2-HG is thought to be mediated by the inhibition of α-KG-dependent dioxygenases, such as the EGLN prolyl-hydroxylases that regulate HIF (15). It is known that acidic pH alone can activate HIF-1α (27), and therefore we hypothesized that 2-HG may play a role in acid-induced HIF activation. To test this, a cell line with stable transgenic over-expression of L-2-HGDH was generated (Figure 8). The L-2-HGDH-Tg cells (clone #2) exhibited 12-fold greater L-2-HGDH activity compared to mock transfected controls (Figure 8B). Furthermore, exposure of cells to acidosis plus KMV as in Figure 1 resulted in elevated 2-HG in control cells with a blunted response in L-2-HGDH-Tg cells (Figure 8C). Measurement of HIF levels by western blot (Figure 9) revealed that HIF stabilization in response to acid plus KMV was significantly blunted in L-2-HGDH-Tg cells (lane 3 vs. 4). In contrast, HIF induction by hypoxia was similar between both control and L-2-HGDH-Tg cells, indicating that canonical HIF signaling was intact. These data indicate that 2-HG generation is required for acid-induced HIF stabilization.

**Figure 8.**
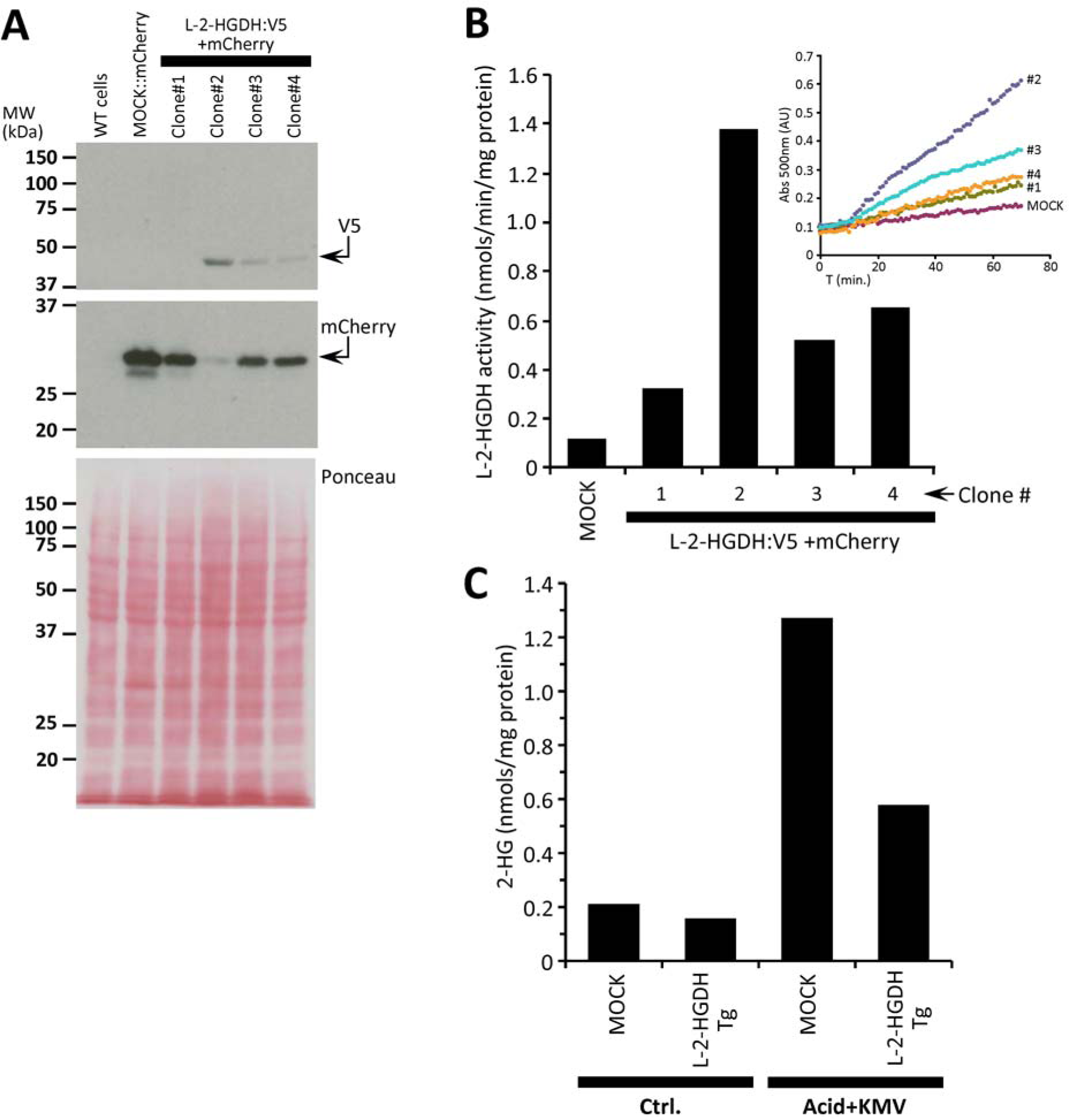
Characterization of L-2-HGDH transgenic cells. HEK293 cells were transfected with mock (mCherry alone) or V5:L-2-HGDH plus mCherry containing vectors, and stable clones selected as described in the methods. **(A):** Monitoring of successful transfection by western blotting for V5 and mCherry. Clone #2 exhibited highest levels of transfection. Lower panel shows Ponceau S stained membrane indicating equal protein loading.**(B):** L-2-HGDH enzymatic activity was measured by the Formazan assay as described in the methods (see also Figure 3 main manuscript). Graph shows calculated rates, with inset showing raw spectrophotometric traces. In agreement with panel A, clone #2 showed highest L-2-HGDH activity. **(C):** Quantitation of 2-HG levels in mock or L-2-HGDH transgenic cells, in response to acid plus KMV exposure as in Figure 1. All data in this Figure are N=1, representing merely rationale for choice of clone #2 for subsequent cell signaling experiments.

**Figure 9.**
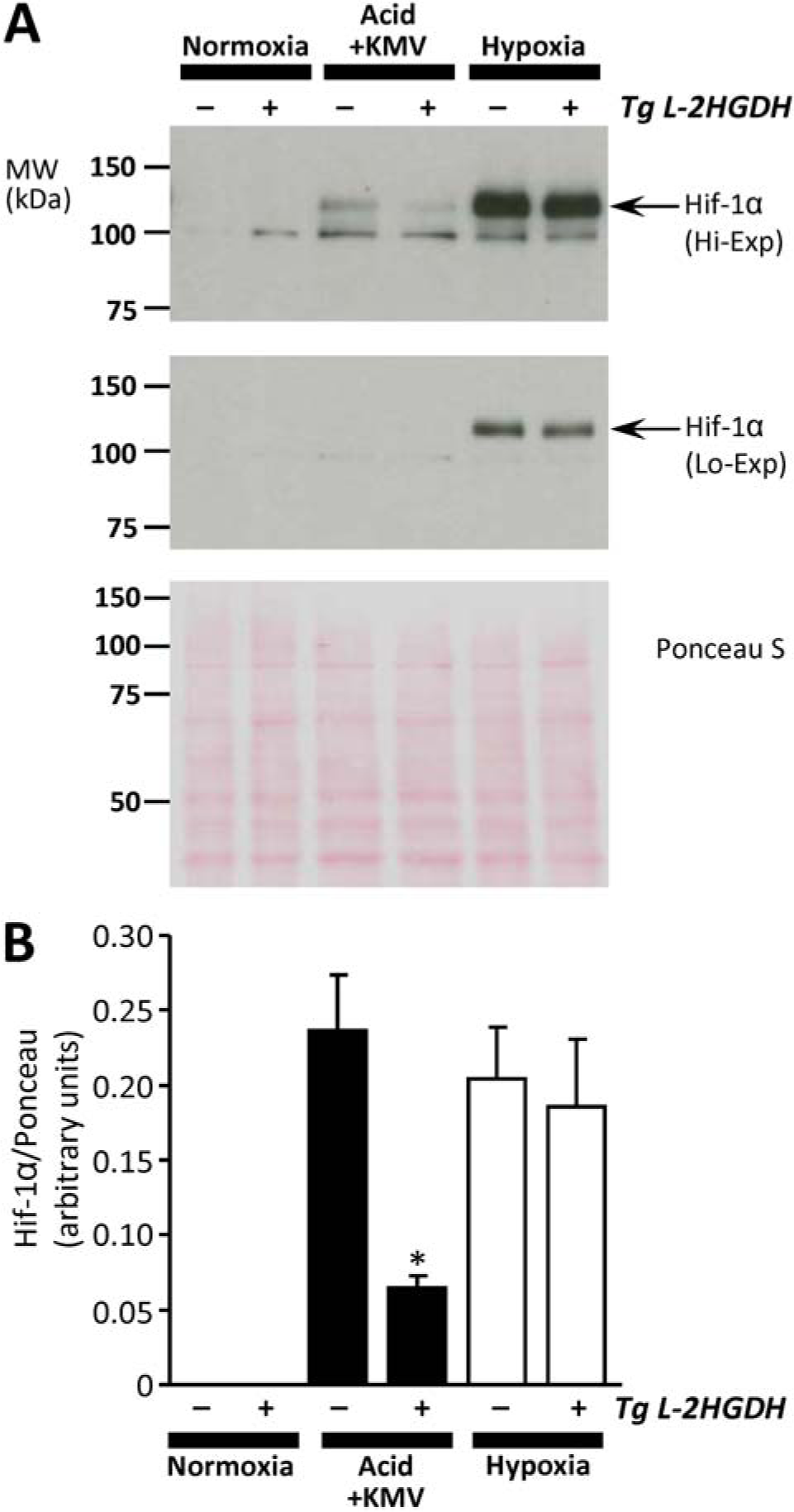
Requirement of 2-HG for acid-induced HIF induction. Mock and L-2-HGDH transfected cells (see Figure 8) were exposed to acid plus KMV as in Figure 1, or to hypoxia, followed by western blotting for HIF-1α. **(A):** Representative HIF-1α western blots at high exposure (upper panel) or low exposure (center panel), with corresponding Ponceau S stained membrane to indicate protein loading (lower panel). Arrows indicate position of HIF-1α at expected molecular weight.**(B):** Quantitative densitometry from 4 independent experiments of the type shown in panel A. For normoxia and acid plus KMV conditions, high exposure blots were used. However, due to saturation of the HIF signal, low exposure blots were used for the hypoxia condition. Data are means ± SEM, N=4. *p<0.05 (paired Student’s t-test) between mock and L-2-HGDH-Tg cells under the same condition.

Although the current data suggest that pH is an important regulator of 2-HG signaling, the relative importance of the pH → 2-HG axis in cancer or stem cell biology remains to be determined. While acidic pH is often associated with the high glycolytic metabolism of cancer cells, the regulation of pH in cancer is complex, with extracellular pH (pH_ex_) tending more toward acidity than intracellular pH (pH_in_) (28, 29). Furthermore, although it has been suggested that exposure to acidic pH_ex_ may play a role in promoting “stem-ness” (30), there has been much recent controversy regarding the effects of acid exposure on stem cell phenotype (31). Thus, while it could be speculated that a simple elevation in 2-HG levels can transmit a pH signal to the epigenetic machinery and bring about a cancer or stem cell phenotype, the situation is likely to be far more complex in-vivo. As the data in Figure 1 suggest, the availability of substrate α-KG is also a major determinant of 2-HG generation, and the fact that α-KG is also a substrate for the dioxygenase signaling enzymes that 2-HG inhibits, suggests that the α-KG/2-HG ratio is likely to be an important determinant of 2-HG signaling outcomes.

Finally, the current findings may also hold therapeutic implications for the hydroxyglutaric acidurias. Lactic acidosis has been reported in some 2-HG aciduric patients (32), and it could be speculated that acidotic episodes may trigger 2-HG accumulation. As such, the careful management of pH to avoid acidosis, for example by treatment with dichloroacetate, (33) may be an overlooked therapeutic approach in these patients.

## EXPERIMENTAL PROCEDURES

All reagents including isolated enzymes were purchased from Sigma (St. Louis, MO) unless otherwise stated. Male C57BL/6J mice were bred in-house from stocks obtained from Jax Labs (Bar Harbor, ME) and were maintained in a thermoneutral environment on a 12 hr. light/dark cycle with food and water available ad libitum. All experiments were in compliance with the NIH “Guide for the Care and Use of Laboratory Animals” and were approved by a local ethics committee (UCAR protocol #2007-087). Enzyme activities were assayed in 25 mM potassium phosphate buffer at 37°C, adjusted to specific pH values as indicated in the figures. Conversion of α-KG to L-2-HG (mediated by LDH, MDH, or ICDH) and was monitored as the oxidation of NADH (0.1 mM) spectrophotometrically at 340 nm. For hypoxia experiments on isolated enzymes, an “open-flow” apparatus was used, in which the spectrophotometer cuvet was fitted with a lid and stirring apparatus, allowing purging of the headspace with gas mixtures defined by mass-flow controllers (34). Gas mixtures were humidified to prevent sample evaporation, and successful achievement of anoxia after 1 hr. of purging with argon was verified by the spectrum of oxy/deoxy-hemoglobin (Hb, 100 μM) in a matching open-flow neighboring cuvet. Oxy-Hb was freshly prepared by dithionite reduction followed by gel filtration and incubation under 100% O_2_ and white light.

The disappearance and formation of L-2-HG, α-KG, citrate/isocitrate and other metabolites was determined by LC-MS/MS analysis of the cuvet reactions. Metabolites were extracted in 80% methanol and then analyzed using reverse phase chromatography with an ion-paring reagent in a Shimadzu HPLC coupled to a Thermo Quantum triple-quadrupole mass spectrometer (7). Data were analyzed using MzRock machine learning tool kit (http://code.google.com/p/mzrock/), which automates analysis of targeted metabolites data based on chromatographic retention time, whole molecule mass, collision energy, and resulting fragment mass. Metabolite concentrations were determined from standard curves constructed using known concentrations.

L-2-HG dehydrogenase activity was assayed (10) in frozen mitochondria isolated from male C57BL6/J mouse hearts as previously described (35). Mitochondria were snap-frozen in liquid N_2_ and stored at −80°C. Mitochondrial protein (0.3 mg/ml) was incubated in 20 mM HEPES buffer supplemented with 0.85 mM MgCl_2_, 0.05 % (vol/vol) Triton X-100, and 1.5 mM iodonitrotetrazolium (ACROS organics). After 5 min., 500 μM L-2-HG was added, and the reaction monitored spectrophotometrically at 500 nm (ε=19300 cm^-1^ M^-1^) for 60 min. At the end of each run, *de-novo* α-KG formation was confirmed by LC-MS/MS.

HEK-293 cells (ATCC, Manassas VA) were maintained in DMEM (Life Technologies, Grand Island NY) with 25 mM D-glucose, 4 mM L-glutamine, 0.1 mM sodium pyruvate, 10 % heat-inactivated FBS (Life Technologies) and 100 µg/ml penicillin-streptomycin (Life Technologies), in 5% CO_2_ at 37°C. For passaging, cells at 75-90% confluence were released by 3 min incubation with 0.25% trypsin-EDTA (Life Technologies), followed by collection in 5 volumes media and pelleting at 500 x *g* for 3 min. Cells were resuspended in 3 volumes media and enumerated on a hemocytometer after Trypan blue staining. Cells were seeded at 10^6^ cells per 75 cm^2^ flask, 24 hr. prior to experiments. Cells were incubated in bicarbonate-free DMEM with 4 mM L-glutamine, 0.1 mM pyruvate, 10 mM glucose, and 10 mM HEPES (adjusted to pH 7.4 or 6.5) at 37°C. Intracellular acidification (19) was achieved by supplementing pH 6.5 media for 15 min. with 30 mM NH_4_Cl (ACROS organics) and 10 µM 5-(N-Ethyl-N-isopropyl) amiloride (EIPA), followed by washing and re-incubation in pH 6.5 media with amiloride alone, for the duration of the experiment (20 hr). This protocol brought intracellular pH to 6.8 within 2 hrs. (Figure 1C). Where appropriate, the α-KGDH inhibitor ketomethylvalerate (KMV) was present at 20 mM. Intracellular pH was monitored by fluorescence microscopy using the ratiometric fluorescent pH indicator 2’,7’-Bis-(2-Carboxyethyl)-5-6-carboxyfluorescein-acetoxymethyl ester (BCECFAM, Invitrogen, Eugene OR) (36). Cells were loaded with 2 µg/ml BCECF-AM) and incubated for 30 min. at 37°C. Cells were then washed once and fluorescence microscopy performed using an Eclipse TE2000-S (Nikon, Avon MA) and data were analyzed using TILL Photonics Imaging System Software. For cellular metabolite analysis, a bolus of 80 % (vol/vol) methanol at −80°C was added after incubations, and cells serially extracted with the same, followed by analysis by LC-MS/MS, as for enzyme assays above.

To create a stable cell line expressing L-2-HGDH, the expression vector pTS4 was created by PCR cloning of the rat L-2-HGDH open reading frame into pcDNA3.1-V5HisC (Invitrogen, Carlsbad, CA) as a Bst*EII* fragment. Rat cDNA was amplified using primers F-5’-ACA CGG TCA CCA TGT GGC CGA CCC TGC GCT ACG-3’ and R-5’-ACA CGG TCA CCT AAC TTA AAC CTT TGC TGT GCT TCC TC-3’. HEK293 cells were transfected with pTFS4 using Lipofectamine 3000 (Invitrogen). Stable clonal isolates were expanded in DMEM with 10 % heat inactivated FBS under 1 mg/ml G418 selective pressure for 10 days and thereafter maintained in media containing 200μg/ml G418. The presence of transgenic L-2-HGDH was monitored by enzymatic assay, and by western blotting for V5 and mCherry tags. pcDNA3.1mCherry was included with pTFS4 at 1:10 to visualize transfected cells, and was transfected on its own to create a negative control (mock transfected) cell line.

For 2-HG signaling experiments, cells were incubated in a hypoxic chamber (<0.1%) at 37 °C for 20 hrs. For western blot monitoring of HIF, cell lysates were harvested under hypoxia in SDS-PAGE sample buffer containing MG132, and following SDS-PAGE proteins were quantified by western blot. Antibodies employed were: anti-V5 (Novex, Carlsbad CA), anti-mCherry (AbCam, Cambridge MA), anti-Hif-1α (Novus Biologics, Littleton CO), anti-β-actin (Santa Cruz Biotech’, Santa Cruz CA). Blots were developed using HRP-linked secondary antibodies and enhanced chemiluminescence, with densitometry in ImageJ software.

All experiments were performed a minimum of 3 times. In purified enzyme studies (Figures 3-5 and Figure 6), “N” represents a single measurement, with multiple measurements made on each of multiple batches of enzyme (i.e. technical replicates, with the number of replicates overall exceeding the number of independent batches of enzyme / biological replicates). In all other Figures, “N” refers to independent biological preparations. Statistical significance between groups was calculated using one-way or two-way ANOVA with post-hoc Student’s t-testing.

## NOTE ADDED IN PROOF

During revision of this manuscript, a report emerged on LDH as a source of high levels of 2-HG found in testis (37), including data showing this phenomenon to be pH sensitive. The relative role of acid as a 2-HG stimulus in testis under physiologically relevant conditions is unclear. We consider this study complimentary to our current findings which span multiple enzymes and cell systems, and overall these discoveries highlight the importance of pH as a driver of metabolic signaling.

## Footnotes

1 Note – The methodology used herein was incapable of distinguishing between L and D 2-HG. However, since the D enantiomer is primarily found in cancers associated with ICDH mutation, while the L enantiomer is primarily made by LDH and MDH under hypoxia, we assume that the bulk of 2-HG measured herein was the L enantiomer.

## ACKNOWLEDGEMENTS

This work was funded by grants from the US National Institutes of Health to PSB (RO1 HL-071158) and JM (R01 AI-081773).

## CONTRIBUTIONS

SMN, JM and XS performed experiments. PSB, SMN, KWN, JM and DF conceived the study. PSB, KWN and JM provided funding and resources. SMN prepared the figures. PSB wrote the paper, with input and editing by KWN, JM and DF.

